# Resequencing and phenotyping of the first highly inbred eggplant multiparent population reveal *SmLBD13* as a key gene associated with root morphology

**DOI:** 10.1101/2025.03.07.642003

**Authors:** Andrea Arrones, Virginia Baraja-Fonseca, Andrea Solana, Mariola Plazas, Salvador Soler, Jaime Prohens, Santiago Vilanova, Pietro Gramazio

## Abstract

The MEGGIC (Magic EGGplant InCanum) population here presented is the first highly inbred eggplant (*Solanum melongena*) multiparent advanced generation intercross (MAGIC) population developed so far, derived from seven cultivated accessions and one wild *S. incanum* from arid regions. The final 325 S5 lines were high-throughput genotyped using low-coverage whole-genome sequencing (lcWGS) at 3X, yielding 293,783 high-quality SNPs after stringent filtering. Principal component analysis (PCA) and neighbour-joining clustering revealed extensive genetic diversity, lack of genetic structure, and the distinct genetic profile of the wild founder. The eight founders and a core subset of 212 lines were phenotyped for above- and below-ground traits, revealing wide phenotypic diversity. Root morphology traits displayed moderate heritability values, and strong correlation were found between root and aerial traits, suggesting that a well-developed root system supports greater above-ground growth. Genome-wide association studies (GWAS) identified a genomic region on chromosome 6 associated with root biomass (RB), total root length (RL), and root surface area (SA). Within this region, *SmLBD13*, a LOB*-*domain protein involved in lateral root development, was identified as a candidate gene. The *S. incanum* haplotype in this region was linked to reduced lateral root branching density, a trait that may enhance deeper soil exploration and resource uptake. These findings provide key insights into root genetics in eggplant, demonstrating MEGGIC potential for high-resolution trait mapping. Furthermore, they highlight the role of exotic wild germplasm in breeding more resilient cultivars and rootstocks with improved root architecture and enhanced nutrient uptake efficiency.

## Introduction

The increasing global population coupled with the rising demand for food and the negative impact of climate change are threatening food security (Yang *et al*., 2024). These challenges underscore the urgent need to achieve a more sustainable and productive horticulture. Addressing these intertwined issues requires the development of innovative and efficient strategies to breed more resilient crops capable of thriving under increasingly unpredictable environmental conditions (Qaim, 2020; Abbas *et al*., 2022). A deeper understanding of complex traits, associated with improved plant performance is crucial, particularly those related to enhanced resource use efficiency, stress tolerance, and overall adaptability to changing climates.

Roots serve as the primary structures anchoring plants to the soil and play an essential role in water and nutrient uptake; therefore, they are key organs affecting plant growth and resilience. Understanding this “hidden half” offers significant potential to optimize crop performance (Maqbool *et al*., 2021). Thus, breeding for more efficient below-ground behaviour might drive a “second green revolution” (Den Heder *et al*., 2010; Uga, 2021). Despite their relevance, root traits have traditionally been neglected in breeding programs due to their phenotyping challenges. Conventional phenotyping methods, such as root digging and soil boring, are labor-intensive and time-consuming. However, advances in phenomic technologies have enabled the development of automated, non-invasive, and high-throughput methodologies (Teramoto and Uga, 2022). Integration of root phenotyping with genomics could further accelerate progress in understanding the genetic basis of root development.

Multiparent advanced generation intercross (MAGIC) populations have emerged as a powerful tool for crop breeding in genetics, ideal for dissecting complex traits (Scott *et al*., 2020). These populations consist of large sets of recombinant inbred lines, offering genetic mosaics of multiple founders suitable for precise fine mapping (Mackay and Powell, 2007; Cavanagh *et al*., 2008; Huang *et al*., 2015). MAGIC populations have been developed in several crops and successfully applied to unravel the genetic basis of different traits of interest, including those related to resilience, such as drought and salt tolerance (Diaz *et al*., 2020; Zhang *et al*., 2020; Abdelraheem *et al*., 2021; Ravelombola *et al*., 2021, 2022; Thudi *et al*., 2023; Sharma *et al*., 2024). To fully exploit the potential of MAGIC populations, deep marker density is required to capture their extensive genetic variation, given their convoluted crossing design and the large population sizes (Arrones *et al*., 2020). Traditionally, two high-throughput genotyping approaches have been predominantly used for MAGIC populations: reduced representation sequencing (RRS)-based methods, such as GBS (López-Malvar *et al*., 2021; Krishnamurthy *et al*., 2022; Sharma *et al*., 2024), and commercial SNP arrays (Yuan *et al*., 2023; Arrones *et al*., 2024; Fourquet *et al*., 2024). While cost-effective, these methods are limited in genome-wide coverage, as they primarily target predefined genomic regions (Barchi *et al*., 2019a; Sun *et al*., 2023). To address this limitation, low-coverage whole-genome sequencing (lcWGS) has emerged as a promising alternative. By combining the broad genomic coverage and dense polymorphism detection of whole-genome sequencing (WGS) with the cost-efficiency of RRS and SNP arrays, lcWGS provides a scalable and cost-effective platform for genome-wide analysis, making it ideal for large-scale studies (Scheben *et al*., 2017; Kumar *et al*., 2021). This approach has been successfully employed in trait-associated loci discovery across various crops, even at ultra-low sequencing coverages as low as 0.02X (Huang *et al*., 2009; Bayer *et al*., 2015; Wang *et al*., 2016; Malmberg *et al*., 2018; Gonda *et al*., 2019; Happ *et al*., 2019; Luo *et el*., 2020; Adhikari *et al*., 2022; Clot *et al*., 2024).

Eggplant (*Solanum melongena* L.) is a major vegetable crop of increasing global significance, ranking fifth in worldwide vegetable production, with an annual output exceeding 60.8 million metric tons in 2023 (FAOSTAT, 2025). Despite its substantial economic and agricultural importance, eggplant has lagged considerably behind other Solanaceae crops, such as tomato, in terms of the development of genetic and genomic resources (Gramazio *et al*., 2017, 2023). To address this gap, we developed the first eggplant MAGIC population, referred to as MEGGIC (Magic EGGplant InCanum). This population was derived from an interspecific cross of seven accessions of cultivated eggplant and one accession of its close wild relative *S. incanum*. The seven *S. melongena* accessions were selected from different geographical origins, including Spain, China, and India, to maximize the phenotypic diversity within the common eggplant, including traits of commercial interest (Gramazio *et al*., 2019; Mangino *et al*., 2022). The *S. incanum* founder was selected to broaden the genetic diversity of the population by introducing ancestral variation lost during domestication processes. The population was advanced to the final stage of five selfing generations (S5) and an optimized lcWGS workflow was developed for genotyping the final S5 MEGGIC lines based on the MEGGIC founders 20X resequencing data (Baraja-Fonseca *et al*., 2024).

Due to root phenotyping difficulties, limited information on genetic control of root development is available for eggplant (Yousefi *et al*., 2024). The wild founder *S. incanum* accession was originally collected from a desertic region in Israel characterized by significant temperature fluctuations between day and night, being exposed to both heat and cold stresses, as well as severe drought conditions (Knapp *et al*., 2013; Gramazio *et al*., 2019). In such environments, a robust root system is critical for plant survival and performance (Delfin *et al*., 2021). A previous study (Flores-Saavedra *et al*., 2024a) demonstrated that genomic introgressions in chromosome 6 from the wild *S. incanum* into cultivated eggplant backgrounds positively influence key traits that enhance overall yield under water stress conditions.

In our study, the final MEGGIC lines were genotyped at 3X lcWGS and screened at the seedling stage for different root morphology traits to identify potential genomic regions associated with an improved eggplant root architecture through genome-wide association studies (GWAS). The identification of lines with improved root systems could represent potential elite material, for direct release as new cultivars, for inclusion in breeding pipelines as pre-breeding resources, or for being used as new rootstocks. Above-ground traits were also evaluated to assess overall plant performance, including some well-known traits in eggplant being used to validate the potential of the MEGGIC population for high-resolution mapping.

We showcased the potential of integrating multiparent populations with low-coverage genomic tools to tackle complex traits such as root architecture, critical for crop adaptation and resilience in the current environmental context. By integrating the assessment of a multiparental MEGGIC population, cost-effective genotyping strategies and high-resolution screening approaches, potential genomic regions associated with improved root traits in eggplant were identified through GWAS. This represents a qualitative leap in eggplant breeding, marking a significant step forward in the development of innovative tools and strategies for genetic improvement.

## Results

### Polymorphisms among MEGGIC lines

The lcWGS genotyping at 3X coverage of the 325 MEGGIC lines yielded 31,673,278 biallelic SNPs with Freebayes v. 1.3.6 (Garrison and Marth, 2012). A final marker set of 293,783 SNPs was selected for subsequent analyses after a rigorous step filtration process (Figure 1.A). The proportion of markers selected from the initial raw set after the comparison with the GS, considered as high-confident biallelic SNPs set as they were supported by at least 20 reads in the GS, was 23.66%, totalling 7,492,731 biallelic SNPs (Figure S1.A). The second filtration step adjusted the SNP heterozygosity of each line, with the proportion of heterozygous SNPs ranging from 0.03 to 0.15 (Figure S1.B). While the minimum depth filtering step increased the proportion of missing data (Figure S1.C and S1.D), this was subsequently addressed in the pre-imputation step by applying a maximum missing data threshold (Figure S1.E and S1.F) and further corrected during the imputation process. The selection of original sites from the fully genotyped dataset resulted in an overall dosage R-squared (DR^2^) of 1, indicating high imputation quality. However, sites with a low allele frequency were removed because of being associated with higher allele-specific error rates (Figure S1.G; Pook *et al*., 2020).

**Figure 1.**
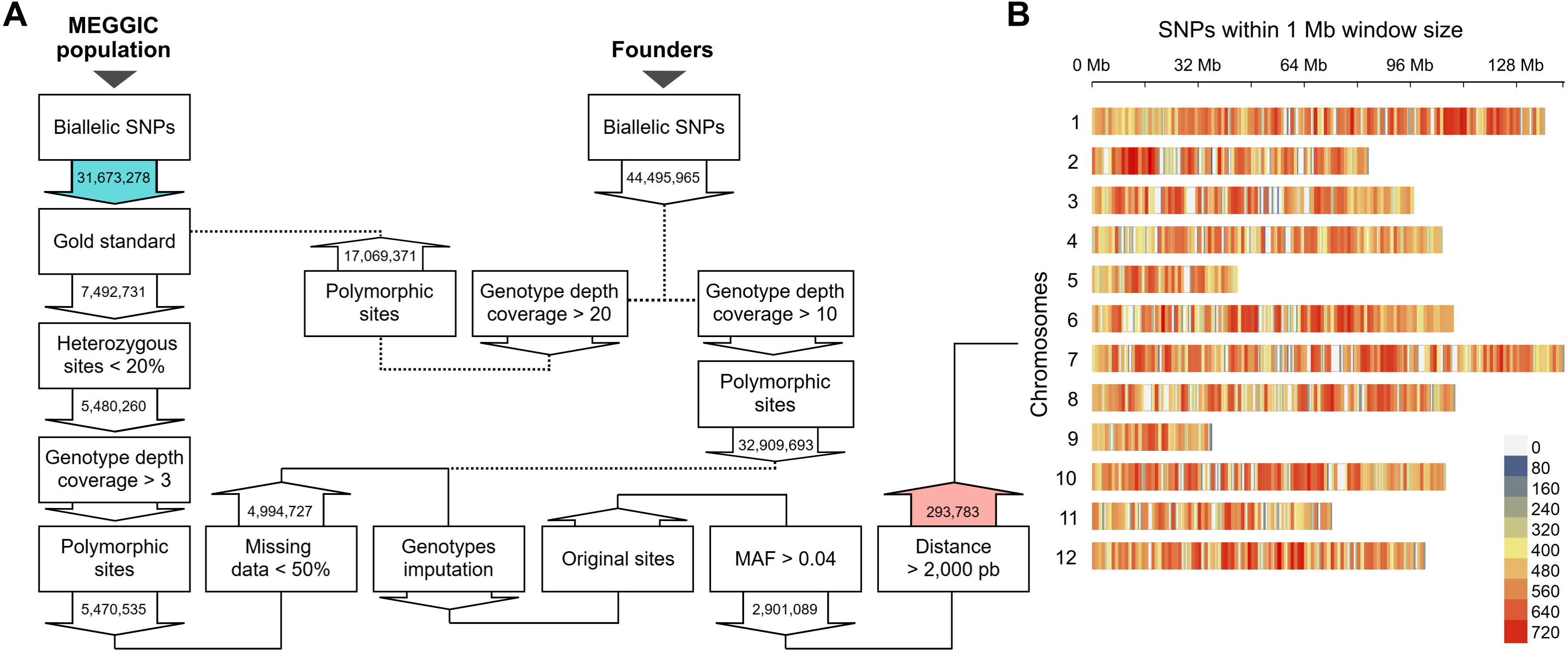
MEGGIC population genotyping results. (A) SNP filtering pipeline from the 3X low-coverage whole-genome sequencing dataset of 31,673,278 biallelic SNPs (blue) to the final subset of 293,783 high-confident SNPs (red) used for the subsequent analysis. The workflow illustrates the impact of each filtering step and the downstream selection of marker subsets. (B) Distribution of the final SNP subset along the 12 eggplant chromosomes. Colour code indicates the SNP density per Mb from lower (blue) to higher (red).

The distribution of SNPs across the 12 eggplant chromosomes was uneven, with the highest number of SNP loci observed on chromosome 1 (38,659 SNPs) and the lowest on chromosome 9 (9,660 SNPs) (Table S1). However, when accounting for physical chromosome length, the SNP density appeared relatively uniform across the genome, with an average of one SNP per 3,931 bp (Figure 1.B, Table S1).

### Population structure and founder contribution

Principal component analysis (PCA) using the genotypic information was performed to evaluate population structure (Diouf and Pascual, 2021). The first principal component (PC1; 5.31%), distinctly separated the wild *S. incanum* from the cultivated *S. melongena* founders, while the PC2 (4.66%) differentiated the Oriental (A and H) from the Occidental (B, D, E, and G) founders, with founder F of unknown origin clustered in the latter group (Figure 2.A; Gramazio *et al*., 2019). Focusing on the main plot where the *S. melongena* parents and MEGGIC lines were distributed, we found that the MEGGIC lines covered a wide area of the zoomed plot with a shift towards positive PC1 values, possibly due to wild introgressions (Figure 2.B). Very importantly, no population structure was detected, as no distinct groups were observed. The first two PCs accounted only for 9.97% of the genetic variance, underscoring the weak population structure and highlighting the high level of genetic diversity. The neighbour-joining dendrogram yielded consistent conclusions (Figure 2.C). The wild founder (C) displayed a highly divergent genetic profile, while the Occidental founders (B, D, E, and G) clustered together reflecting their closer genetic relationship. Specifically, founders D and E, as well as founders B and G, tightly clustered in the same branch indicating a strong genetic similarity between these pairs.

**Figure 2.**
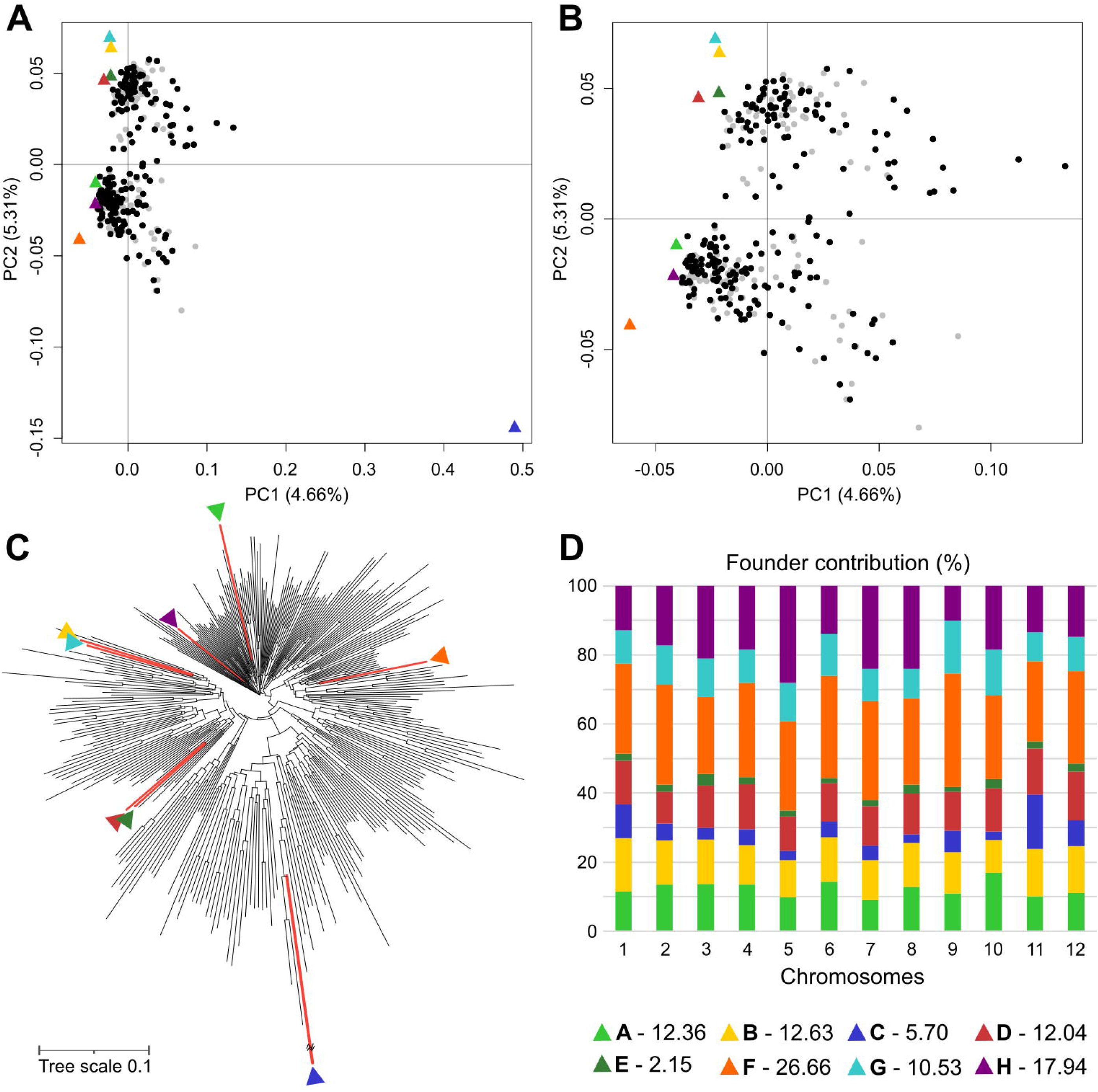
MEGGIC population structure analysis. (A) Principal component analysis (PCA) including the genotyping data of the founders and the MEGGIC population, highlighting the set of lines selected for this study (black dots) over the entire population (grey dots). Founders are indicated with triangles using a colour code. (B) Zoom in on the top left-hand corner of the PCA. (C) Dendrogram indicating founders’ location with coloured red branches and triangles with the colour code. (D) Chromosome-wide and genome-wide allele contribution of parental lines. Numbers in the legend indicate the overall genome-wide contribution of each parental line.

The reconstruction of the genomic mosaics in the MEGGIC lines, based on the eight founder haplotypes, revealed differential haplotype block proportions across different genomic regions for all chromosomes (Figure S2). The eight-founder crossing design theoretically predicts an equal contribution of approximately 12.50% from each founder to the genetic diversity of the final population. However, genome-wide and chromosome-wide assessments of parental allelic probabilities revealed deviations from this expectation, with estimated genomic contributions varying across chromosomes. On average, founder contributions align more closely with the expected value than those reported for the S3MEGGIC at an intermediate stage of the population development (Mangino *et al*., 2022). However, some founders contributed disproportionately to the estimated genetic background of the population, with founders F and H showing the highest estimated average contributions at 26.66% and 17.94%, respectively. In contrast, founders C and E contributed the least to the estimated contribution to the genetic background, with averages of 5.70% and 2.15%, respectively (Figure 2.D).

### Morphological phenotyping

Nine traits, four related to the aerial growth and development (AB, HE, LN, and LA) and five related to root morphology (RB, RL, SA, MD, and MW), were assessed across MEGGIC founders at the seedling stage. Initial evaluation of founder genotypes revealed significant diversity across all measured traits (Figure 3). Notably, the wild *S. incanum* founder (C) exhibited a distinctive aerial morphology and root architecture compared to the other founders. In this way, the aerial part of this founder was marked by higher plant height and leaf number although coupled with lower leaf expansion. In contrast, the root system was characterized by minimal lateral root development but significantly deeper roots (Figure S3). The substantial diversity observed among founders prompted further investigation into plant performance across the entire population to unravel the genetic basis underlying these traits. Consequently, the MEGGIC population was evaluated under the same experimental conditions and following the same methodology. Additionally, PR and AN traits were also collected. A core subset of MEGGIC lines was selected based on seed availability and germination consistency, including lines with data from at least two reliable replicates. After data curation, a core subset comprising 212 lines was retained for further analysis, which was representative of the genetic diversity observed in the entire population (represented as black dots over grey dots in Figure 2.B). These remaining 212 MEGGIC lines displayed a broad range of variation highlighting the wide phenotypic diversity within the population (Table 1). Most traits displayed a continuous distribution slightly skewed to the right (Figure 4.A). Aerial traits exhibited heritability values ranging from 0.21 to 0.31. In contrast, root morphology traits showed higher heritability values ranging from 0.29 to 0.51.

**Figure 3.**
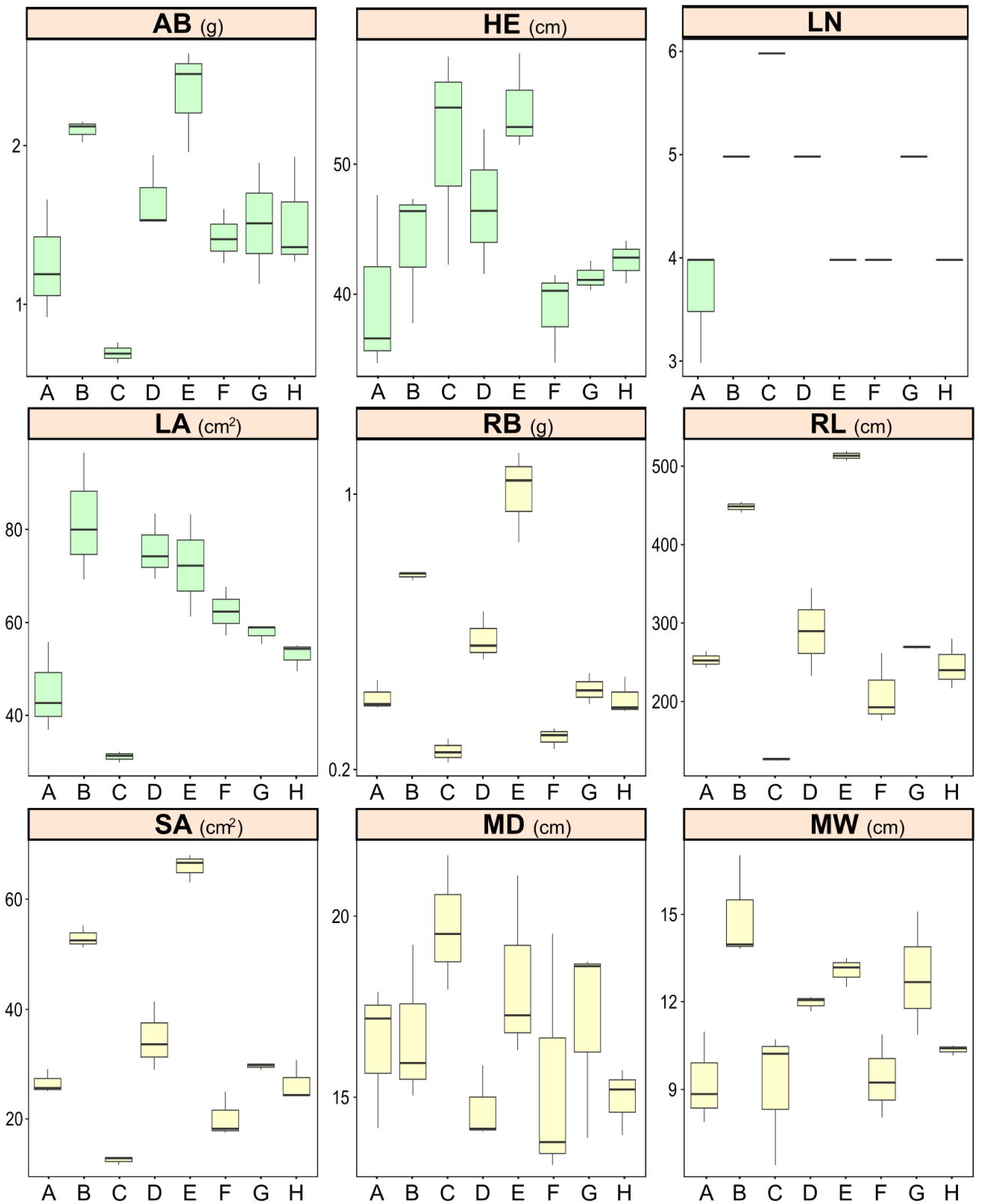
Boxplots illustrating the diversity among MEGGIC founders (A-H) for different aerial (in green: aerial biomass, AB; plant height, HE; leaf number, LN; and leaf area, LA) and root (in yellow: root biomass, RB; total root length, RL; surface area, SA; maximum depth, MD; and maximum width, MW) traits.

**Figure 4.**
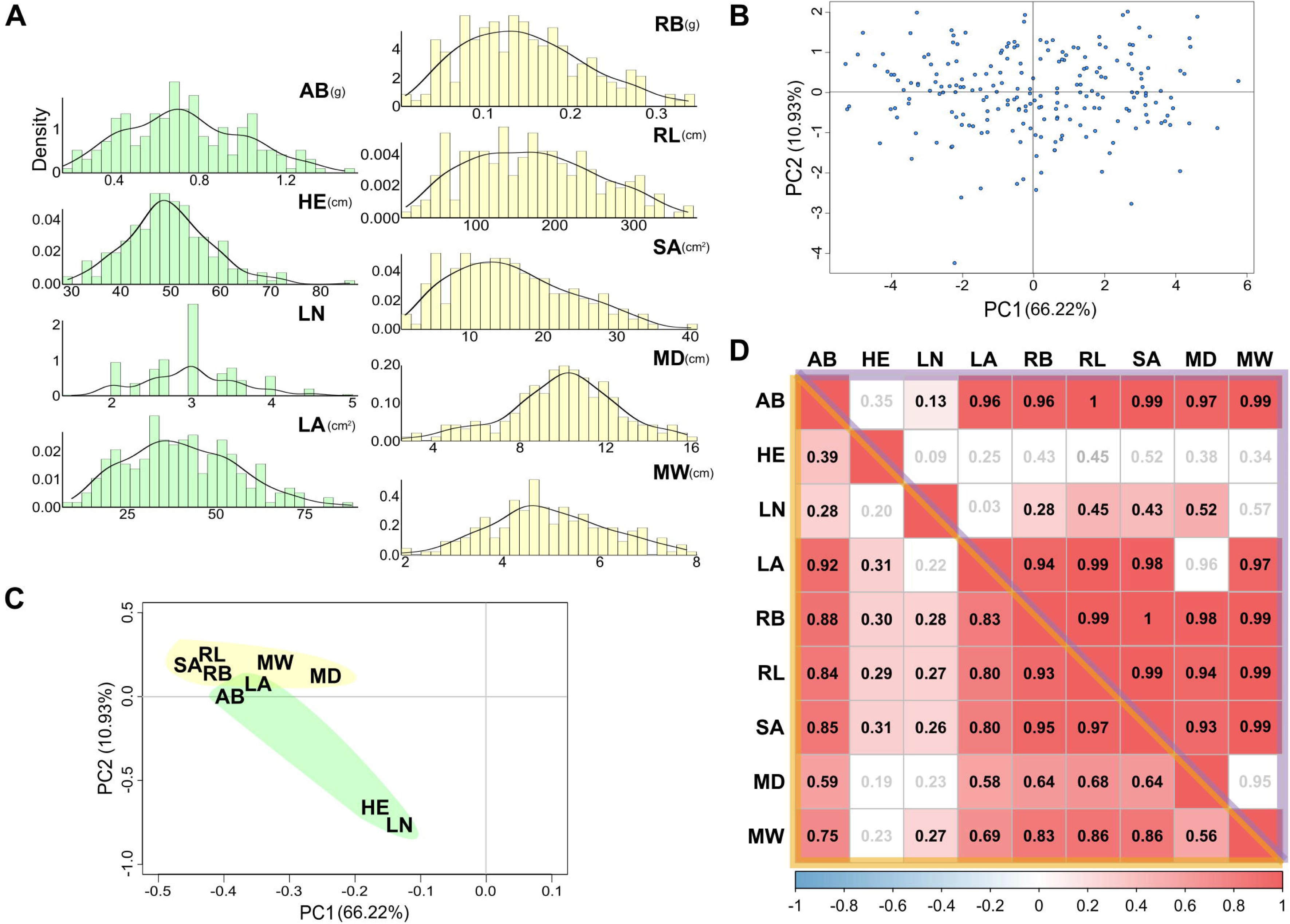
Statistical analysis of the MEGGIC seedlings phenotyping. (A) Histograms and density plots of the phenotype values among MEGGIC core subset lines for different aerial (in green: aerial biomass, AB; plant height, HE; leaf number, LN; and leaf area, LA) and root (in yellow: root biomass, RB; total root length, RL; surface area, SA; maximum depth, MD; and maximum width, MW) traits. (B) PCA score plot and (C) loading plot on the first principal components based on all the studied traits for the selected MEGGIC lines. (D) Pairwise phenotypic (orange lower-left matrix) and genetic (purple upper-right matrix) correlations among the studied traits. Pearson’s correlation coefficient (r) is shown using a Bonferroni correction at the significance level of 0.05. Blue and red colours correspond to negative and positive correlations, respectively.

**Table 1.** Mean (± standard deviation), range values, genetic variance 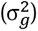, residual variance 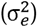, and boad-sense heritability (H^2^) with standard errors (within brackets) of the different traits evaluated in the MEGGIC core subset lines.

| Trait | MEGGIC lines (n=212) |  |  |  |  |
| --- | --- | --- | --- | --- | --- |
| | Mean | Range | $\sigma_g^2$ | $\sigma_e^2$ | $H^2$ |
| *Aerial growth* |  |  |  |  |  |
| AB (g) | 0.73 ± 0.36 | 0.08-2.25 | 0.03 | 0.09 | 0.24 (0.049) |
| HE (cm) | 49.49 ± 11.03 | 24.93-92.82 | 36.54 | 82.26 | 0.31 (0.051) |
| LN | 2.98 ± 0.78 | 1-6 | 0.16 | 0.44 | 0.27 (0.050) |
| LA (cm<sup>2</sup>) | 40.69 ± 20.72 | 6.13-139.18 | 82.67 | 318.89 | 0.21 (0.047) |
| *Root morphology* |  |  |  |  |  |
| RB (g) | 0.15 ± 0.09 | 0.01-0.60 | 0.002 | 0.006 | 0.29 (0.050) |
| RL (cm) | 171.21 ± 95.60 | 9.96-593.76 | 4,862.84 | 4,647.16 | 0.51 (0.047) |
| SA (cm<sup>2</sup>) | 15.66 ± 9.61 | 0.85-65.13 | 40.88 | 51.83 | 0.44 (0.050) |
| MD (cm) | 9.96 ± 3.19 | 2.01-19.48 | 3.13 | 6.86 | 0.31 (0.051) |
| MW (cm) | 4.96 ± 1.50 | 1.36-10.74 | 0.65 | 1.54 | 0.30 (0.051) |

The phenotypic PCA performed with the evaluated traits in the MEGGIC lines facilitated the assessment of variation across multiple traits within the population and revealed the relationship among lines. The first two PCs accounted for 66.22% and 10.93% of the total variation, respectively. The projection of the lines in the PCA score plot showed a wide distribution over the graph area, which underscored a substantial phenotypic diversity within the MEGGIC lines (Figure 4.B). Regarding the PCA loading plot, all the traits were placed on the negative values of PC1 (Figure 4.C). Aerial-related traits were mainly located in or close to the negative values of PC2, while root morphology-related traits were plotted close together in the positive values of PC2.

Pearson’s correlation coefficients were calculated to estimate phenotypic correlations both within and across aerial and root traits (Figure 4.D). For the aerial traits, only AB and LA exhibited a high positive correlation (0.92), suggesting that LA is a good indicator of the plant’s overall growth potential. For root morphology traits, strong positive correlations were observed among all traits, with correlations ranging from 0.56 to 0.97 (mean of 0.79). RL and SA showed a high correlation of (0.97), and both also showed a strong correlation with RB (0.93 and 0.95, respectively), all of them contributing to the overall efficiency of the root system. There was a positive correlation between root traits, AB, and LA with values ranging from 0.58 to 0.88 (mean of 0.76), suggesting that a robust root system supports greater above-ground growth, especially in terms of leaf expansion. Genetic correlations were estimated and found to closely align with the phenotypic correlations (Figure 4.D).

By analysing MEGGIC lines distribution for each trait, several lines were identified that exhibited high vigour in both above- and below-ground development (i.e. MEGGIC lines 101 or 148), indicating strong overall growth potential (Figure S4). Conversely, some lines were observed to display significantly weaker development in both aerial and root systems (i.e. lines 102 or 210). Some lines were found to prioritize LA expansion over LN (i.e. line 114) or prioritize LN production over root development (i.e. line 73).

### GWAS analysis and candidate gene identification

To validate the suitability of the MEGGIC population for GWAS, PR and AN traits were initially analysed, since they are well-characterized and associated QTLs and genes have been proposed. Given the absence of population structure, a GLM model was conducted. The Manhattan plot for PR revealed a strong association peak at the end of chromosome 6 with the highest significantly associated SNP located at 105,615,359 bp, explaining 36.81% of the total phenotypic variance (Figure 5.A). This peak is very close to the *SmLOG3* gene (SMEL_006g267050.1, 105,504,136-105,509,693 bp), which has been recently identified as a major gene for prickliness in eggplant (Satterlee *et al*., 2024). The Manhattan plot for AN displayed a pronounced association peak on chromosome 9, with the top significantly associated SNP positioned at 17,400,739 bp, accounting for 25.61% of the overall phenotypic variance (Figure 5.B). Near this SNP, the *SmbHLH69* gene (SMEL_009g326640, 17,862,102-17,872,412 bp), also known as *SmTT8*, was identified, which promotes anthocyanin biosynthesis in eggplant (He *et al*., 2019; Shi *et al*., 2021).

**Figure 5.**
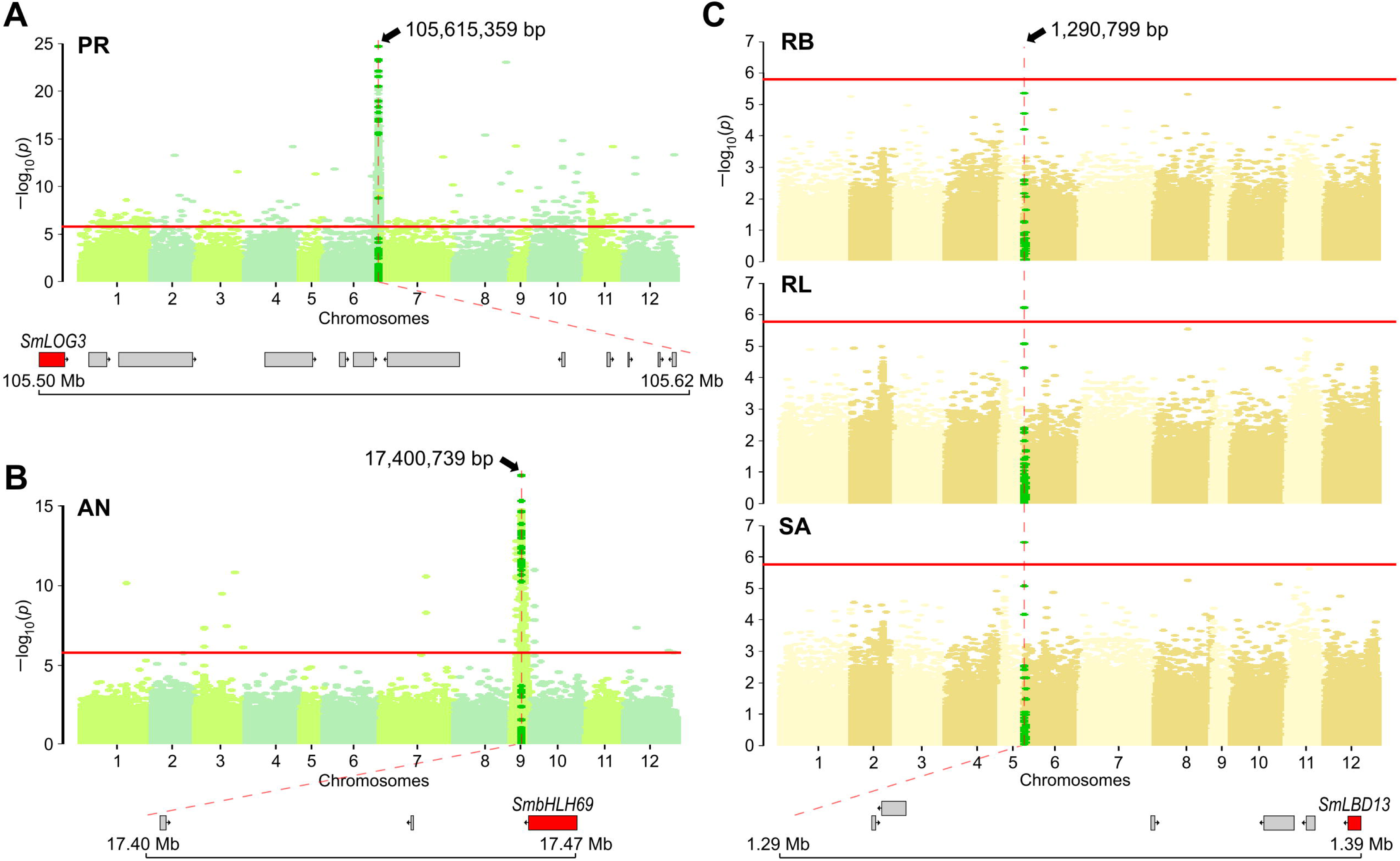
Genome-wide association studies (GWAS) results for (A) prickles (PR), (B) anthocyanin pigmentation (AN), and (C) different root-related traits, including root biomass (RB), total root length (RL), and surface area (SA). The horizontal red lines represent the Bonferroni threshold at p = 0.05 (LOD = 5.77). The vertical dashed red line indicates the associated genomic position. Bright green dots covered the neighbouring positions to the top significantly associated SNP. Genes close to the top significantly associated SNP position are shown, with the proposed candidate gene indicated in red.

For root-related traits, GWAS analyses were performed to investigate genetic control underlying root development. The Manhattan plot for RL and SA revealed a single peak on chromosome 6, with the SNP over the Bonferroni threshold (LOD = 5.77) located at 1,290,799 bp, explaining 12.33% of the total phenotypic variance (Figure 5.C). Although no SNP exceeded the Bonferroni threshold for RB, there was a clear upward trend at the same position. No trend was observed for MD and MW traits (Figure S5). Under the peak on chromosome 6, the *SmLBD13* gene (SMEL_ 006g243020.1, 1,386,770-1,388,917 bp) was identified as the best candidate (Table S2). This gene belongs to the lateral organ boundaries (LOB) domain-containing proteins, which are known to play crucial roles in plant development (Xu *et al*., 2016). Variants that predicted high-impact effects on protein function were annotated by SnpEff for founders C, D, and H. Specifically, for founders C and H, a SNP (C/A) leading to a stop gain was identified in two different positions (at 1,388,151 and 1,388,725 bp, respectively); while for founder D a frameshift variant (TTGT/TT) was identified at 1,387,319 bp. These variants were supported by a limited number of reads, which may compromise their reliability (Table S3). For this reason, a comparative analysis of the founder haplotype diversity was performed for the associated genomic region where *SmLBD13* is located. As a result, it was observed that MEGGIC lines carrying the founder C haplotype displayed on average reduced lateral root branching density, contributing to smaller RB, RL, and SA mean values (Table 2). In contrast, lines carrying the founder B haplotype exhibited on average a more fibrous root system. No assessed lines were found to carry the founder E haplotype in this genomic region, consistent with its overall low estimated contribution to the population for this region.

**Table 2.** Mean (± standard deviation) of the root-related traits evaluated in the MEGGIC core subset lines based on the founders’ haplotype in the associated genomic region on chromosome 6.

| Founder | MEGGIC lines' haplotype average |  |  |
| --- | --- | --- | --- |
|  | RB (g) | RL (cm) | SA (cm<sup>2</sup>) |
| A | 0.13 ± 0.08 | 155.72 ± 84.36 | 14.64 ± 8.70 |
| B | 0.22 ± 0.13 | 220.87 ± 138.11 | 22.25 ± 13.48 |
| C | 0.10 ± 0.06 | 112.83 ± 69.43 | 10.45 ± 6.68 |
| D | 0.13 ± 0.09 | 135.93 ± 93.48 | 12.73 ± 8.92 |
| E | - | - | - |
| F | 0.14 ± 0.06 | 162.61 ± 69.57 | 14.60 ± 6.70 |
| G | 0.17 ± 0.07 | 207.79 ± 79.90 | 19.00 ± 7.53 |
| H | 0.14 ± 0.06 | 165.56 ± 66.73 | 14.90 ± 6.96 |

## Discussion

A comprehensive understanding of traits governing crop adaptation and resilience is crucial to cope with increasingly unpredictable environmental conditions. Among these traits, root architecture plays a pivotal role in enhancing the plant ability to adapt to environmental stresses, improving overall plant performance and resource use efficiency (Wang *et al*., 2019; Katuuramu *et al*., 2020, Yousefi *et al*., 2024). In this context, MAGIC populations have emerged as a cutting-edge resource due to their potential for high-precision association studies, establishing it as a valuable resource for genetic mapping and trait dissection. In this study, we presented the final version of the first MAGIC population in eggplant (MEGGIC) constituted by 325 highly inbred lines. Among the eight founders, founder C corresponds to an accession of the close wild relative *S. incanum* (Mangino *et al*., 2022), whose genetic distinctiveness provides the opportunity to introgress unique traits into a cultivated background (Flores-Saavedra *et al*., 2024b).

lcWGS provides a cost-effective platform for genome-wide analysis, making it particularly suitable for large-scale studies (Han *et al*., 2020; Fang *et al*., 2023; Thudi *et al*., 2023). Here, following Baraja-Fonseca *et al*. (2024) recommendations, the 325 MEGGIC lines were high-throughput genotyped at 3X coverage, which provided accuracy, sensitivity, and genotypic concordance comparable to 5X coverage while significantly reducing sequencing costs. However, low sequencing coverages inherently pose challenges due to the limited number of reads per site, potentially leading to erroneous variant calls (Meisner and Albrechtsen, 2018). To address this drawback, rigorous SNP filtering is critical, with the implementation of a GS being a widely recognized approach for ensuring genotypic reliability (Happ *et al*., 2019; Gao *et al*., 2021; Zook *et al*., 2020). In the context of inbreed populations, GSs are typically constructed using high-coverage resequencing data from the founders (Diaz *et al*., 2020; Wang *et al*., 2022; Saripalli *et al*., 2023; Clot *et al*., 2024). Despite the limited number of individuals included, these GSs capture the full allelic diversity of the population, as no novel alleles are expected beyond those present in the founder genomes. In this study, we implemented a filtering step strategy involving the comparison of the dataset against a GS and the application of a minimum depth coverage threshold (Baraja-Fonseca *et al*., 2024) to minimize potential false positives and generate a high-confidence set of biallelic SNPs with reliability comparable to variants surveyed at 20X coverage. This genotyping strategy resulted in a significantly higher SNP density (one SNP per 4 kb) and a more evenly distributed set of markers compared to the single-primer enrichment technology (SPET) (Barchi *et al*., 2019a) used for the genotyping of the intermediate-stage S3MEGGIC population, which achieved a density of one SNP per 165 kb (Mangino *et al*., 2022).

Using the genotyping data, the genetic diversity and population structure of the MEGGIC population were thoroughly analysed. A PCA including the founders’ genetic information reflected the genetic divergence of the wild founder, which was clearly separated from the cultivated founders and the rest of the lines in the population. Genetic similarities were observed within the cultivated founders, as evidenced by their clustering in both the PCA and dendrogram analyses, particularly among those of Occidental origin. The wide distribution of the MEGGIC lines in the PCA graph area, coupled with the low variance explained by the first PCs, highlighted the extensive genetic diversity within the population and the lack of population structure, which is one of the main advantages of this kind of multiparent populations (Mackay and Powel, 2007). A deeper genotyping characterization of the population enabled a more precise reconstruction of haplotypes, providing more accurate estimations of founder haplotype proportions and better capturing the fine-scale genomic structure of the population (Pook *et al*., 2020). However, genome-wide and chromosome-wide analyses revealed deviations from the expected equal contribution of each founder from an 8-way crossing design, a pattern also observed in other MAGIC populations (Hashemi *et al*., 2022). Despite an increased marker density, the genetic similarity among some founders may hinder the reliable distinction between them, potentially introducing bias in the estimation of haplotype blocks. The lower contribution of the wild founder could also be attributed to segregation distortion and cryptic selection processes due to reduced fertility, recalcitrant germination, or erratic flowering and fruit set, since they have previously been reported in progenies from crosses between *S. incanum* and cultivated eggplant (Lefebvre *et al*., 2002; Barchi *et al*., 2010).

The 212 MEGGIC lines phenotyped covered a broad genetic diversity of the entire population indicating that this core subset of lines was well-suited for mapping QTL or genes of interest. Seedlings were grown in expanded clay balls which facilitates straightforward root extraction and non-destructive root extraction, enabling high-quality root scanning. This growing system provides an efficient approach for root phenotyping and could be extended to other crops to facilitate studies on root architecture and development. Seedlings were phenotyped for different aerial and root-related traits showing a wide range of variation. Moderate heritability values were obtained for root morphology traits, suggesting that selection for improved root architecture, could lead to significant and consistent genetic gains (Mathew and Shimelis, 2022; Schuster *et al*., 2024). Since positive correlations were observed among root and aerial traits, selecting for improved root traits could indirectly enhance above-ground growth, promoting overall plant vigour and resilience under stress conditions (Zhang *et al*., 2024). Moreover, positive correlations between traits exhibiting differing heritabilities suggested that an indirect selection approach by targeting a genetically correlated trait with higher heritability may represent a more effective strategy for improving traits with lower heritability, thereby optimizing breeding efforts (Neyhart *et al*., 2019). Some MEGGIC lines exhibited different biological trade-offs between above-and below-ground resource allocation, underscoring diverse adaptation strategies for balancing growth and resource use (Weigelt *et al*., 2021).

As proof of concept for testing the potential of the MEGGIC population for the high-precision fine mapping of traits of interest, two extensively studied traits in eggplant, PR and AN, were selected. GWAS analysis for PR directly pointed to the well-known *prickleless* (*pl*) locus on chromosome 6 (Frary *et al*., 2014), very close to *SmLOG3*, recently identified as the responsible gene for prickle losses across the *Solanum* as well as in distantly related vascular plant lineages (Satterlee *et al*., 2024). For AN, a differential trend was observed in the genetic regulation across developmental stages of the MEGGIC population. In the intermediate-stage S3MEGGIC population, GWAS for for anthocyanin presence in vegetative plant tissues and fruit epidermis identified a major peak on chromosome 10 near to the *SmMYB113* gene, a well-known regulatory transcription factor controlling anthocyanin synthesis in eggplant (Mangino *et al*., 2022). Additionally, a peak on chromosome 9 was detected close to *SmTT8*, although with a lower LOD score. In contrast, here in the more advanced S5 MEGGIC population, the strongest association signal for anthocyanin presence was located on chromosome 9, near to the *SmTT8* gene. The key difference between the S3MEGGIC and the final MEGGIC phenotyping was the plant development stage (adult plants vs 25-day-old seedlings, respectively). Given that *SmTT8* is a basic-helix-loop-helix (bHLH) protein that directly binds to *SmMYB113* via their amino acid terminus domain to module anthocyanin biosynthesis (Barchi *et al*., 2019b; Moglia *et al*., 2020; Zhou *et al*., 2020; Shi *et al*., 2021), these results suggest a shift in the genetic regulation of anthocyanin accumulation with stage-specific expression patterns. The *SmTT8* gene seems to be predominantly expressed during the early stages of plant development, with increased activity in vegetative tissues, while expression may shift towards a higher activation of *SmMYB113* gene during reproductive development and in fruit tissues (Petroni and Tonelli, 2011). Further expression analysis at different developmental stages would provide valuable insights into the dynamic interaction between *bHLH* and *MYB* genes in anthocyanin regulation.

Understanding the variation in root traits and identifying candidate SNPs or genes associated with this variation is essential for breeding eggplant root systems. Different root-related traits were selected for further GWAS analysis. Since there was a strong correlation among traits (r > 0.93), similar Manhattan plots were obtained. As a result, an associated genomic region was identified at the beginning of chromosome 6. The candidate genomic region colocalized with the *SmLBD13* gene (SMEL_ 006g243020.1), which belongs to lateral organ boundaries (LOB) domain-containing proteins. These proteins are known to be key regulators of a large number of developmental and metabolic processes in higher plants, particularly in lateral organ formation, including lateral root development and root architecture plasticity (Xu *et al*., 2016). LOB domain proteins are involved in regulating processes like lateral root initiation and formation in other species, such as rice (Liu *et al*., 2005), maize (Taramino *et al*., 2007), or *Arabidopsis* (Cho *et al*., 2019). Identifying this gene as a key regulator of root morphology in eggplant has important implications for understanding root architecture, which could enhance breeding efforts aimed at improving nutrient uptake efficiency, stress resilience, and overall plant performance. The ‘steep, cheap, and deep’ root ideotype for rapid exploration of deeper soil layers, improving water and nutrient uptake, is crucial for optimizing plant plasticity (Lynch, 2022). One of the key mechanisms contributing to this ideotype is the reduction of lateral root branching density. The presence of the wild haplotype in the associated genomic region contributed to a less fibrous root system in the MEGGIC lines. These results were consistent with the *S. incanum* root morphology shaped by the harsh and resource-scarce environments where it evolved (Knapp *et al*., 2013). However, given that dense root hairs are important for the acquisition of immobile soil nutrients, especially phosphorus and potassium (Lynch, 2022), lines carrying the *S. melongena* founder B haplotype, characterized by a more fibrous root system, may also hold agronomic value. The identification of promising lines that combine a more efficient root system provides valuable candidates for further analysis as potential elite breeding materials and might also be useful as rootstocks for eggplant grafting.

Our study directly addresses global challenges related to population growth, climate change, and the urgent need for sustainable agriculture by providing genetic insights that contribute to the development of more resilient crops (Yang *et al*., 2024). The highly inbred MEGGIC population represents a milestone in eggplant research, as it integrates high-throughput genotyping with phenomic-based phenotyping to unravel the genetic control of root architecture. Through high-resolution genetic mapping, we identified key genomic regions associated with root system traits, highlighting *SmLBD13* as a strong candidate gene for lateral root development. These findings provide valuable genetic resources for breeding strategies aimed at enhancing eggplant adaptability to increasingly unpredictable environments. By leveraging the power of MAGIC populations and cost-effective genomic tools, this study marks a significant step forward in improving root traits, a critical yet often overlooked aspect of crop resilience.

## Experimental procedure

### MEGGIC population development and genotyping

#### Plant materials

The MEGGIC (Magic EGGplant InCanum) population was developed by intercrossing seven cultivated common eggplant (*S. melongena*; A, B, D, E, F, G, and H) and one wild relative (*S. incanum*; C). Founders were pairwise intercrossed by following a simple “funnel” approach (Figure S6). After obtaining four simple hybrids (AB, CD, EF, and GH) and two double hybrids (ABCD and EFGH), the latter were intercrossed following a chain pollination scheme obtaining 209 combinations of quadruple hybrids (S0 generation) (Mangino *et al*., 2022). Two plants of each S0 progeny were randomly selected and selfed during five generations (S5) by single seed descent (SSD). Due to the loss of some lines during the SSD process and to some plants failing to set fruit or produce seed, the final MEGGIC population is constituted of 325 highly inbred lines.

#### Genotyping of the MEGGIC population

The 325 MEGGIC lines were germinated in seedling trays. Young leaf tissue was sampled from each line and genomic DNA was extracted using the silica matrix extraction (SILEX) method (Vilanova *et al*., 2020). The quality and integrity of the extracted DNA were assessed by agarose gel electrophoresis and NanoDrop spectrophotometer ratios (260/280 and 260/230), while its concentration was estimated using a Qubit 2.0 Fluorometer (Thermo Fisher Scientific, Waltham, MA, United States). All samples were high-throughput genotyped by lcWGS at 3X (3.6 Gb clean data each) using the DNBseq platform at Beijing Genomics Institute (BGI Genomics, Hong Kong, China) following Baraja-Fonseca *et al*. (2024) recommendations. Initial raw reads were processed for quality control using fastq-mcf v.1.04.676 (Aronesty, 2013). High-quality reads were aligned onto the eggplant reference genome “67/3” v. 3 (Barchi *et al*., 2019b), using the BWA-MEM algorithm v. 0.7.17–r1188 (Li, 2013) with default parameters. PCR duplicates were removed by using MarkDuplicates software from Picard’s tools v. 1.119 (https://broadinstitute.github.io/picard/).

#### SNP selection

Variant detection was performed with Freebayes v. 1.3.6 (Garrison and Marth, 2012), using the standard settings except for a minimum quality threshold of 20 for both mapping and base quality. Biallelic SNPs were selected using BCFtools v. 1.13 (Li, 2011). To filter out potential false positives from lcWGS, the dataset was benchmarked against a high-confidence reference set (hereafter gold standard, GS) developed by Baraja-Fonseca *et al*. (2024). This set, generated from 20X resequencing data of the MEGGIC population founders (Gramazio *et al*., 2019), consists of biallelic SNPs supported by a minimum of 20 reads. Shared positions between the 325 MEGGIC cohort and the GS underwent a rigorous four-step filtration process: (I) elimination of positions with over 20% heterozygosity among lines; (II) conversion of genotypes supported by fewer than three reads to missing data; (III) removal of monomorphic sites across lines; and (IV) removal of sites with more than 50% missing data. For complete genomic representation, imputation was carried out with Beagle v. 22Jul22.46e (Browning and Browning, 2016), using the GS as the reference panel, with additional filtering for a minimum depth of 10 using VCFtools v. 0.1.16 (--minDP 10). Only original positions with a minor allele frequency greater than 0.04 was retained, maintaining a minimum distance of 2,000 bp between selected positions to ensure data quality and reduce redundancy.

#### Population structure and founder contribution

A principal components analysis (PCA) was conducted to study the pattern of genetic variation among the final MEGGIC lines based on the filtered SNPs. Founders’ genetic information was also included. The genetic matrix was calculated using the prcomp function from the stats package in R (R Core Team, 2023). The eigenvalues of each principal component (PC) and the proportion of explained variance were used to evaluate the structure of the MEGGIC population. The two first PCs were drawn using R package ggplot2 (Wickham, 2016). A dendrogram was constructed using the neighbour-joining method (Saitou and Nei, 1987), and the graphical representation was generated and refined using iTOL v.4 software (Letunic and Bork, 2019). To estimate the parental contribution to the final population, haplotype blocks, and recombination patterns within the population, the R package HaploBlocker v. 1.7.01 was used (Pook *et al*., 2019).

### MEGGIC population phenotyping and analysis

#### Seedling cultivation conditions and experimental design

In an initial exploratory trial, the eight founders were evaluated for trait diversity, followed by assessing the entire population for traits that exhibited significant variation. Seeds of the eight founders and the 325 MEGGIC lines were germinated in Petri dishes, following the Ranil *et al*. (2015) protocol to overcome the seed dormancy commonly observed in wild species and synchronized germination. Subsequently, they were transferred to seedling trays of 0.2 l containing expanded clay pebbles of 2–3 mm diameter (Arlita™, Madrid, Spain) using a completely randomized block design with three replicates. Seedlings (one per replicate) were grown in a climatic chamber under a photoperiod and temperature regime of 16 h light (25 °C, 100–112 μmol m^−2^ s^−1^) and 8 h dark (18 °C) and 70% of relative humidity. They were cultivated for 25 days, and they were fertirrigated three times a week with 50 ml of ¼ strength Hoagland no. 2 solution (Hoagland and Arnon, 1950).

#### Phenotyping of the MEGGIC seedlings

After 25 days of cultivation, seedlings were cut and partitioned into aerial and root parts for the evaluation of four traits related to the aerial growth and development (aerial biomass, AB; plant height, HE; leaf number, LN; and leaf area, LA) and five traits related to roots morphology (root biomass, RB; total root length, RL; surface area, SA; maximum depth, MD; and maximum width, MW) as indicated in Table 3. Additionally, the presence of prickles (PR) and anthocyanin pigmentation (AN) in the leaves and stem were assessed for validating MEGGIC potential for high-precision fine mapping. The aerial part was photographed with a 4 cm^2^ red calibration area for LA analysis with the Easy Leaf Area software (Figure S7) (Easlon and Bloom, 2014). Default parameters were used except for the minimum red-green-blue (RGB) values and G/B ratio which were set at 0 and 2, respectively. Roots were spread in a transparent tray in a thin layer of water, and they were imaged using a high-resolution scanner. Image acquisition was performed by WinRhizo root scanner (dual lens system STD 4,800 root scanner Epson Perfection V700, Regents Instrument Canada Inc.) and analysis was carried out by RhizoVision Explorer (Figure S7) (Seethepalli *et al*., 2021). Non-root objects were filtered at a maximum size of 2 units and root pruning was enabled with a threshold set at 5 units. MD and MW were measured by Image J software (Abràmoff *et al*., 2004).

**Table 3.** List of traits used for the MEGGIC characterization with their abbreviations, units, and method of collection.

| Abbreviation | Trait | Units | Method of collection |
| --- | --- | --- | --- |
| *Aerial characteristics* |  |  |  |
| AB | Aerial biomass | g | Weighed using a precision balance |
| HE | Plant height | cm | Measured from ground level to apical meristem |
| LN | Leaf number |  | Including mature and young leaves |
| LA | Leaf area | cm<sup>2</sup> | Easy Leaf Area software (Easlon and Bloom, 2014) |
| PR | Prickles |  | Binary classification for absence (0) or presence (1) |
| AN | Anthocyanins |  | Measured from none (0) to very dark pigmentation (3) |
| *Root morphology* |  |  |  |
| RB | Root biomass | g | Weighed using a precision balance |
| RL | Total root length | cm | RhizoVision Explorer (Seethepalli *et al.*, 2021) |
| SA | Surface area | cm<sup>2</sup> | RhizoVision Explorer (Seethepalli *et al.*, 2021) |
| MD | Maximum depth | cm | Image J software (Abràmoff *et al.*, 2004) |
| MW | Maximum width | cm | Image J software (Abràmoff *et al.*, 2004) |

#### Statistical analysis

For each trait, the phenotypic mean and range values were calculated for the three replicates. To visualize the distribution of these traits, histograms and density plots were generated using the R package ggplot2 (Wickham, 2016). Principal Component Analysis (PCA) was performed using the prcomp function to assess the phenotypic variation among the MEGGIC lines, with score and loading plots created using ggplot2. Pearson’s correlation coefficients between traits were calculated and significance was assessed with a Bonferroni correction at a significance level of 0.05 (Pearson, 1895; Hochberg, 1988), using the R packages psych (Revelle, 2017) and corrplot (Wei *et al*., 2017). Additionally, barplots of standardized mean values with error bars were produced using ggplot2 to identify interesting MEGGIC lines.

Genomic heritabilities (H²) were estimated by fitting univariate linear mixed models using the genomic best linear unbiased prediction (GBLUP) framework (Clark and van der Werf, 2013). Fixed effects only included the intercept and MEGGIC lines were treated as random genetic effects, assumed to be normally distributed with mean 0 and variance equal to genomic relationship matrix (GRM). Residuals were assumed to be independently and normally distributed. The models were fitted using the R package sommer (Covarrubias-Pazaran, 2016). The genomic relationship matrix was constructed following VanRaden (2008) method 1 using R package AGHmatrix (Amadeu, 2016). The genomic heritability for each trait was calculated using the formula

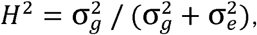

where 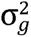 is the genetic variance using marker-based relationships and 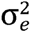 is the residual variance. Standard errors for the heritabilities were obtained using the vpredict function in sommer. Additionally, genetic correlations between traits were estimated by fitting a multivariate linear mixed model using the GBLUP framework within sommer. Fixed effects included only the intercept and an unstructured variance model between traits was used for both the random genetic effects and residuals to allow correlations across traits to be estimated. Genetic correlations were calculated in R using cor(rho_pheno[lower.tri(rho_pheno)], rho_geno[upper.tri(rho_geno).

#### Genome-wide association study (GWAS)

PR and AN traits were used for validating the potential of the MEGGIC population for GWAS analysis since these traits have been extensively studied in eggplant (Barchi *et al*., 2012; Cericola *et al*., 2014; Frary *et al*., 2014; Toppino *et al*., 2020; Mangino *et al*., 2022; Satterlee *et al*., 2024). Several quantitative trait loci (QTLs) and candidate genes have been proposed to control these traits, so we aimed to assess whether the results obtained in the MEGGIC population were consistent with the previously reported regions. Additionally, five root-related traits were analysed as key indicators of root development since they are suggested to be directly related to nutrient acquisition and crop yield (Wang *et al*., 2019; Katuuramu *et al*., 2020, Yousefi *et al*., 2024). The phenotyping data as the mean value of the three replicates for each trait was used and GWAS analyses were performed using the TASSEL software (ver. 5.0, Bradbury *et al*., 2007). General linear model (GLM) analysis was conducted for the association study (Price *et al*., 2006). The multiple testing was corrected with the Bonferroni method (Holm, 1979) with a significance level of 0.05 (Thissen *et al*., 2002). SNPs with a limit of detection (LOD) score (calculated as -log10[p-value]) exceeding these specified thresholds or cutoff values in both GWAS models were considered significantly associated with the traits under evaluation. The genes surrounding the highest significant SNPs were retrieved from the “67/3” v. 3 eggplant reference genome (Barchi *et al*., 2019b). Candidate genes were assessed by SnpEff prediction software v 4.2 (Cingolani *et al*., 2012) based on resequencing data from the eight MEGGIC founders (Gramazio *et al*., 2019) to identify causative mutations associated with phenotypic variation. The Integrative Genomics Viewer (IGV) tool was then used to visually explore the founder genome sequences and validate the SnpEff predictions (Robinson *et al*., 2023). Founder haplotypes were estimated for the candidate genomic region and a comparative analysis of founder haplotype diversity across the MEGGIC lines combining genotypic and phenotypic data was performed.

## Acknowledgements

This work has been funded by grants PID2021-128148OB-I00 funded by MICIU/AEI/10.13039/501100011033/ and by ERDF/EU, PDC2022-133513-I00 funded by MICIU/AEI/10.13039/501100011033/ and European Union Next Generation EU/PRTR, CIPROM/2021/020 from Conselleria d’Educació, Cultura, Universitats i Ocupació (Generalitat Valenciana), and by the Horizon Europe programme, project number 101094738 (“Promoting a Plant Genetic Resource Community for Europe; PRO-GRACE). VB-F is grateful to Conselleria d’Educació, Cultura, Universitats i Ocupació (Generalitat Valenciana), for a predoctoral contract (CIACIF/2023/238). AS is grateful to MICIU/AEI/10.13039/501100011033/ and FSE+ for a predoctoral grant (PRE2022-102368). Pietro Gramazio is grateful for the post-doctoral grant RYC2021-031999-I funded by MICIU/AEI/10.13039/501100011033 and the European Union through NextGenerationEU/PRTR.

## Author contributions

MP, SS, JP, SV, and PG conceived the idea and supervised the manuscript. AA and AS performed the root phenotyping trials. AA, VB-F, PG, and SV performed the bioinformatic analysis and analyzed the data. AA prepared a first draft of the manuscript. All other authors reviewed and edited the manuscript.

## Data availability statement

The datasets presented in this study can be found in online repositories. The names of the repository/repositories and accession number(s) can be found below: https://www.ncbi.nlm.nih.gov/, PRJNA392603 and PRJNA1174391.

