## Supplementary figures and images for "Resequencing and phenotyping of the first highly inbred eggplant multiparent population reveal *SmLBD13* as a key gene associated with root morphology"

### Figure S1

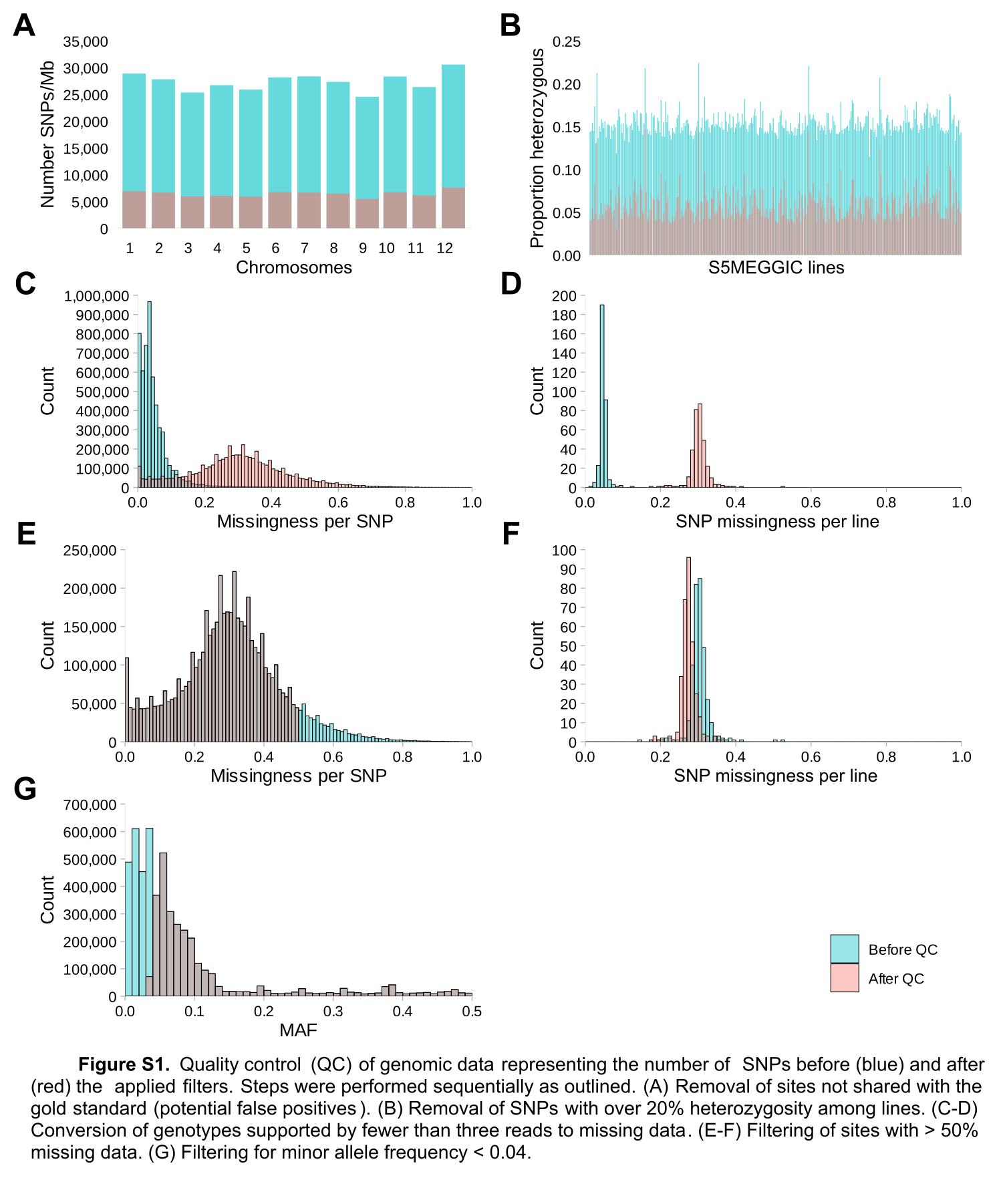

### Figure S2

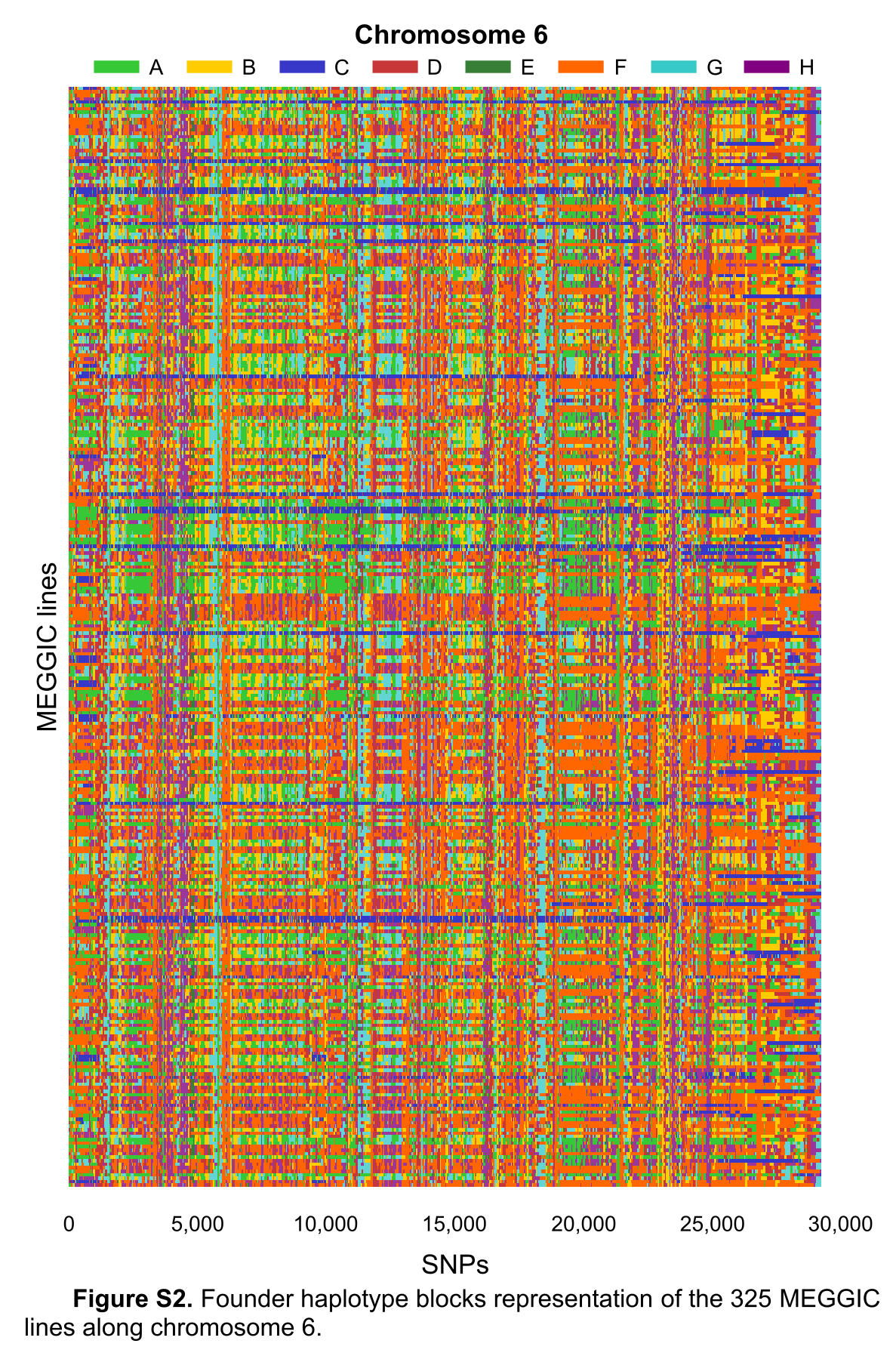

### Figure S3

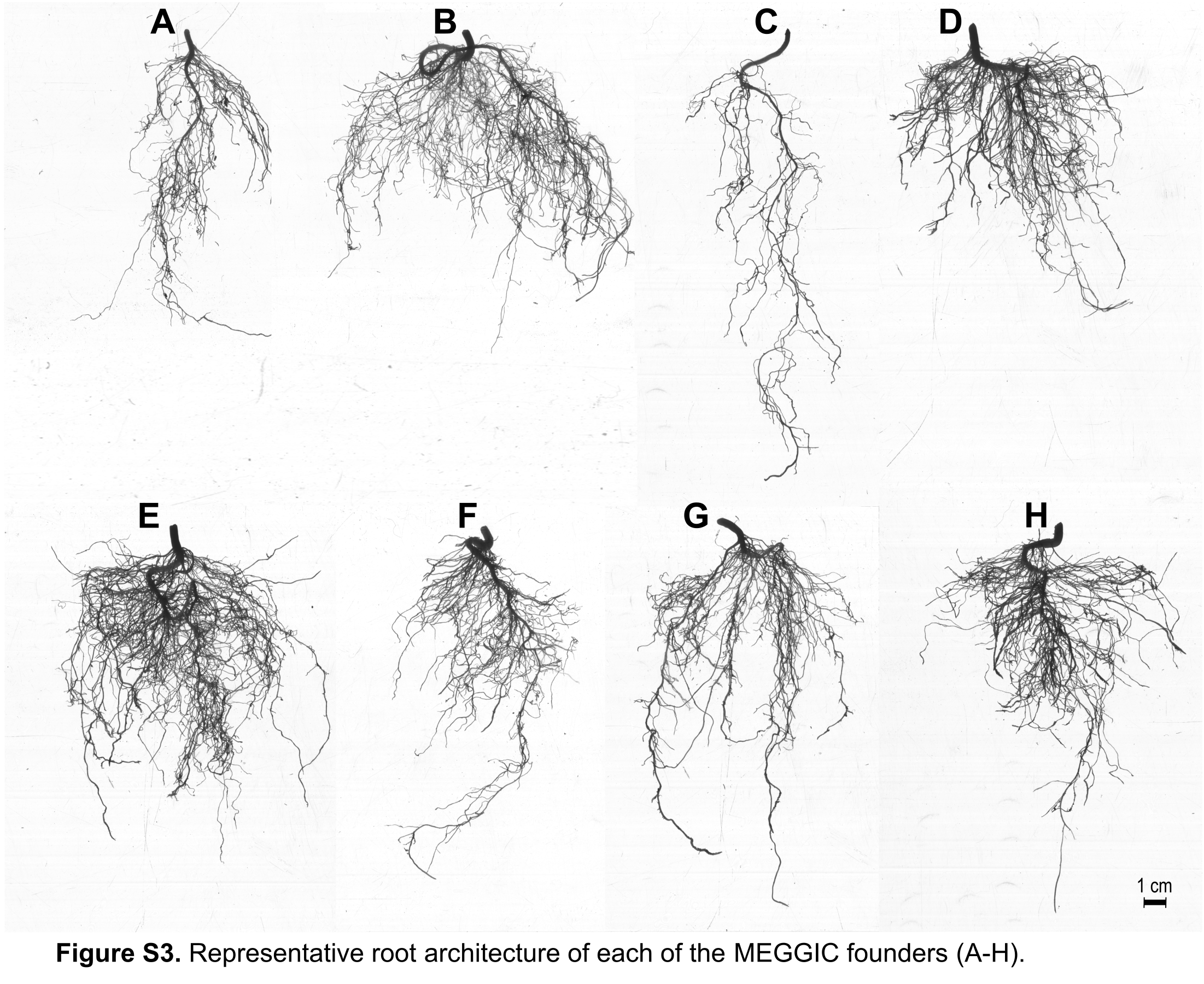

### Figure S4

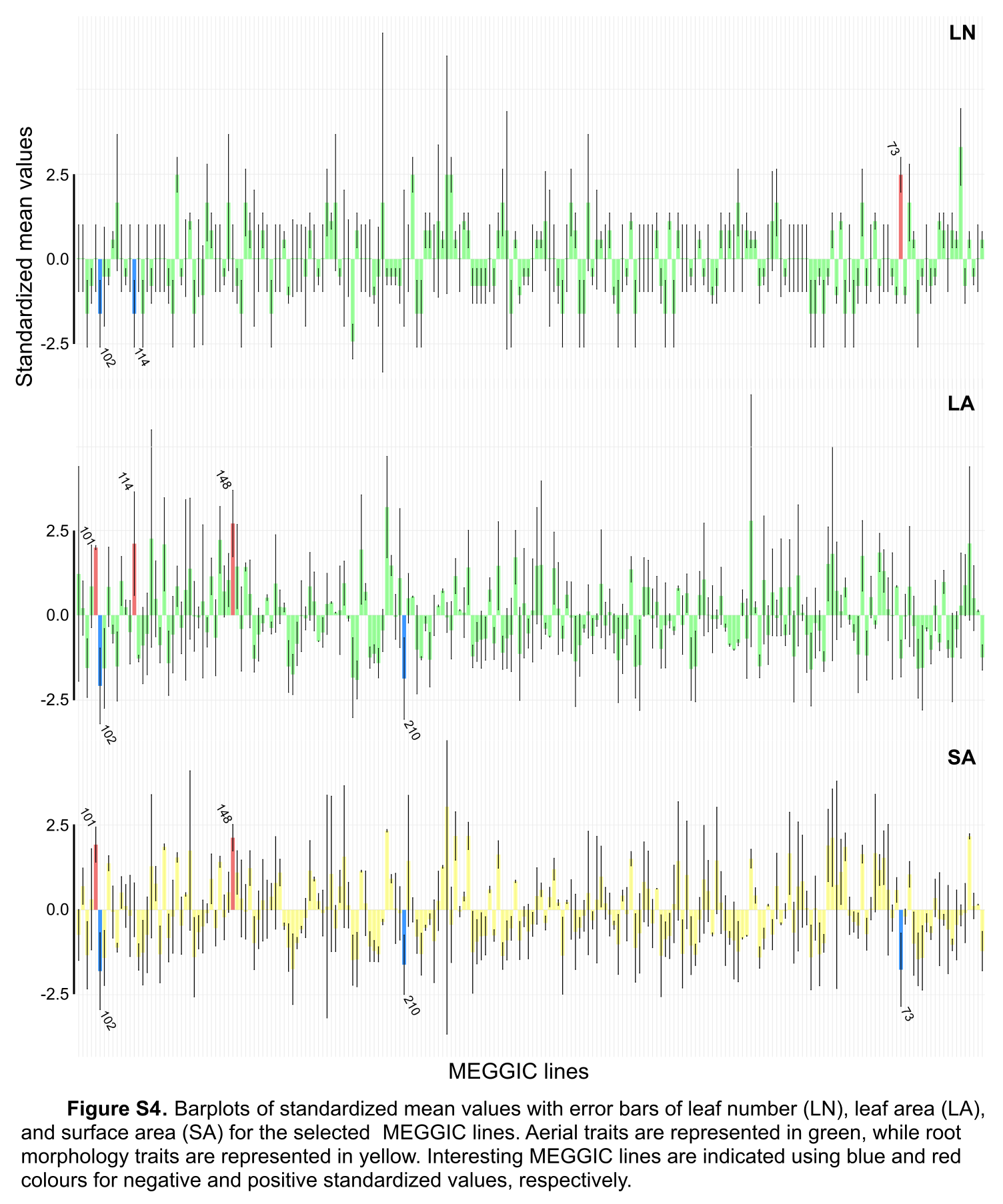

### Figure S5

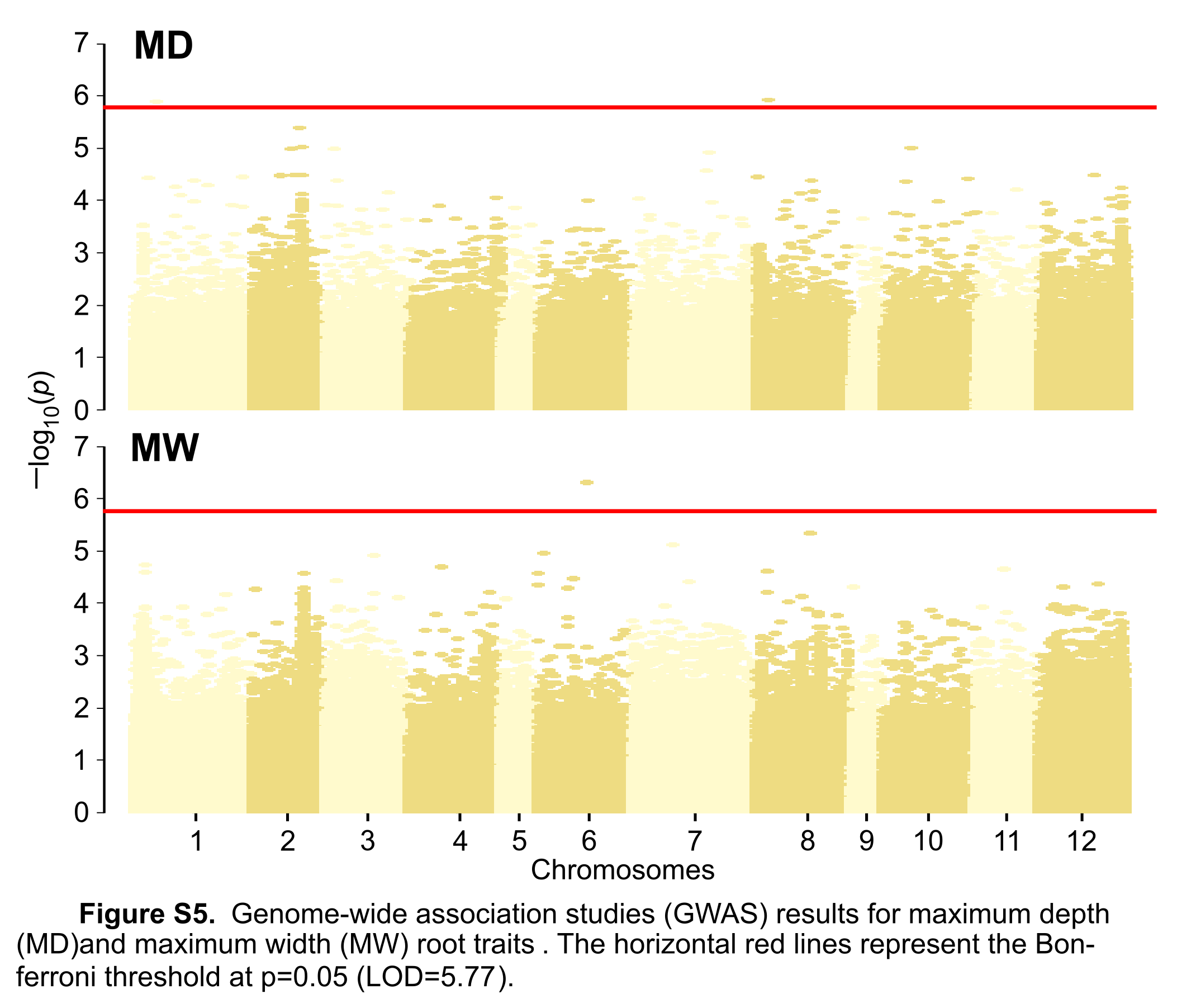

### Figure S6

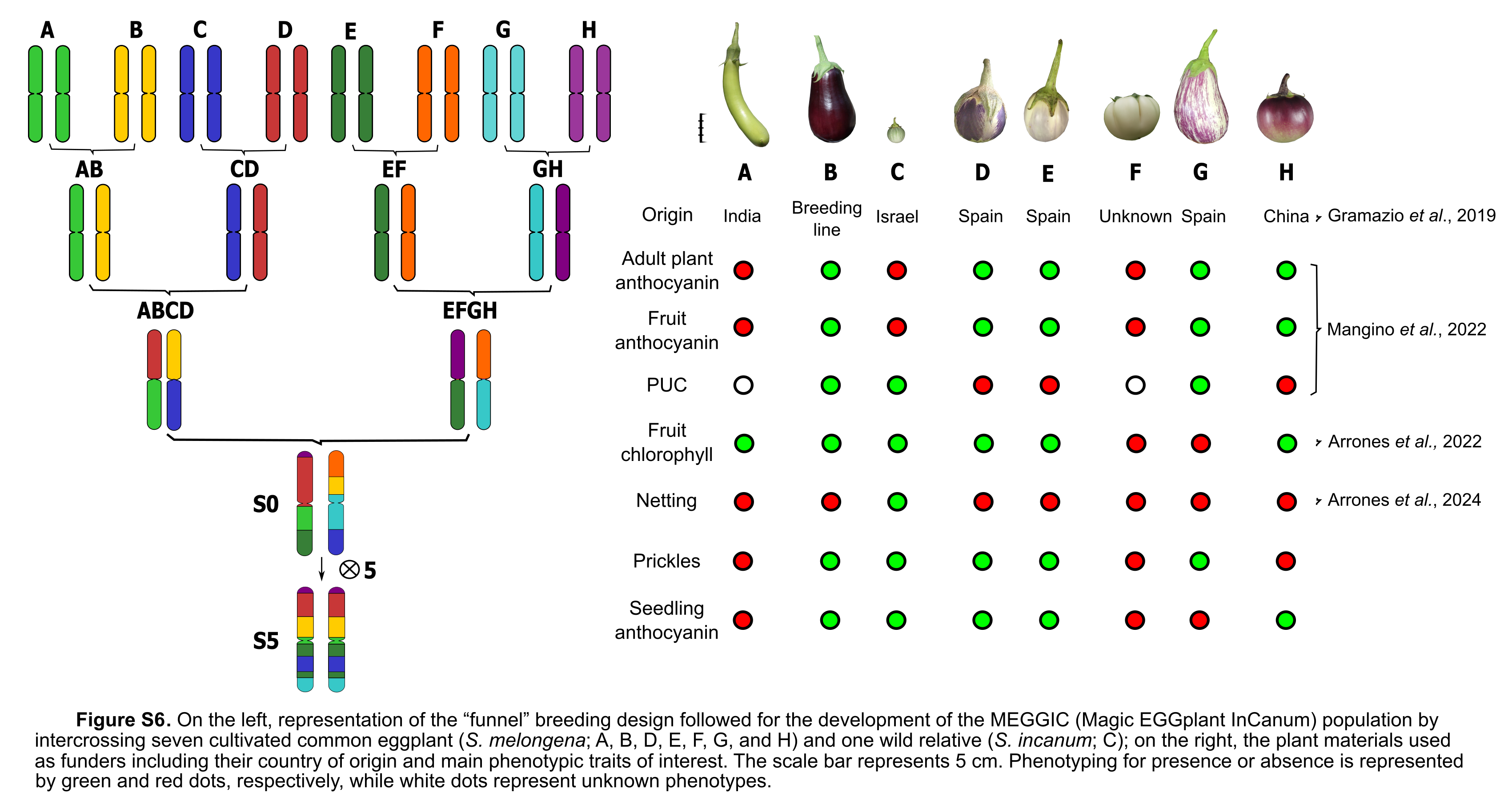

### Figure S7

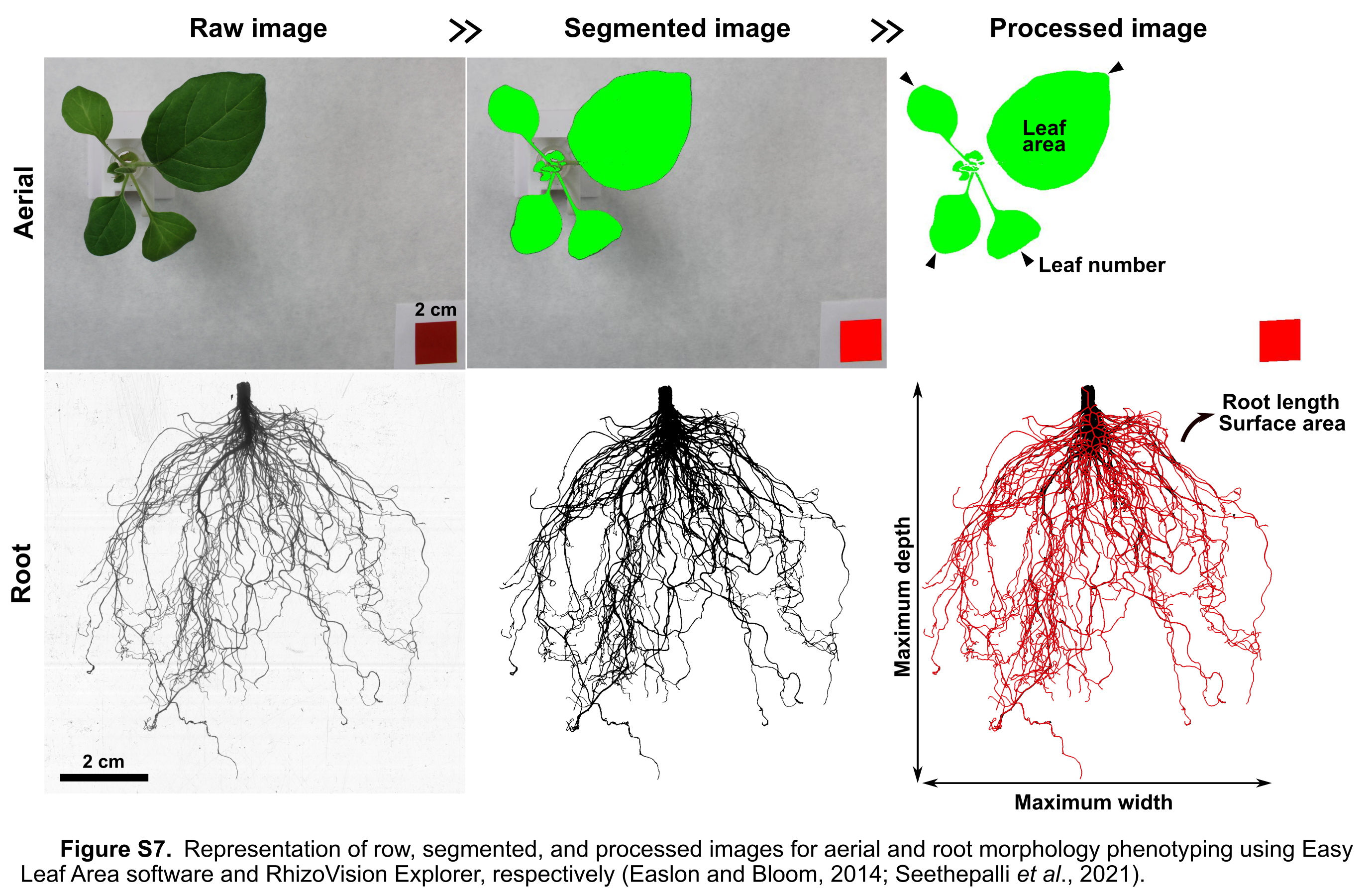
